# Temporal properties of painful contrast enhancement using repetitive stimulation

**DOI:** 10.1101/2021.08.12.456139

**Authors:** Tibor M. Szikszay, Waclaw M. Adamczyk, Juliette L. M. Lévénez, Philip Gouverneur, Kerstin Luedtke

**Affiliations:** Institute of Health Sciences, Department of Physiotherapy, Pain and Exercise Research Luebeck (P.E.R.L.), University of Luebeck, Luebeck, Germany; Laboratory of Pain Research, Institute of Physiotherapy and Health Sciences, The Jerzy Kukuczka Academy of Physical Education, Katowice, Poland; Institute of Medical Informatics, University of Luebeck, Luebeck, Germany

## Abstract

Offset analgesia is characterized by a disproportionately large reduction in pain following a small decrease in a heat stimulus and is based on the phenomenon of temporal pain contrast enhancement (TPCE). The aim of this study is to investigate whether this phenomenon can also be induced by repetitive stimulation, i.e., by stimuli that are clearly separated in time. With this aim, the repetitive TPCE paradigm was induced in healthy, pain-free subjects (n=33) at the volar non-dominant forearm using heat stimuli. This paradigm was performed applying three different interstimulus intervals (ISIs): 5, 15, and 25 seconds. All paradigms were contrasted with a control paradigm without temperature change. Participants continuously rated the perceived pain intensity. In addition, electrodermal activity was recorded as a surrogate measure of autonomic arousal. Temporal pain contrast enhancement was confirmed for both ISI 5 seconds (p < 0.001) and ISI 15 seconds (p = 0.005), but not for ISI 25 seconds (p = 0.07), however the magnitude of TPCE did not differ between ISIs (p = 0.11). Electrodermal activity was consistent previous pain ratings, but showing significantly higher autonomic activity being measured. Thus, the phenomenon of temporal contrast enhancement of pain can also be induced by repetitive stimulation. Both the involvement of the autonomic nervous system and the involvement of habituation processes are conceivable, which consequently points to both central and peripheral mechanisms of TPCE.

**Summary:** The temporal contrast enhancement of pain and electrodermal activity can be provoked by stimuli that are clearly separated in time.

## 1. Introduction

Acute and chronic pain perception depend on an interaction between peripheral noxious afferents and the processing in the central nervous system [6]. Pain perception can be influenced by faciliatory (pro-nociceptive) and inhibitory (anti-nociceptive) aspects of endogenous descending pain modulation [60] and may therefore be a main contributing factor in the development and maintenance of persistent pain [2,36,59]. One frequently used paradigm reflecting those aspects of anti-nociceptive pain modulation is offset analgesia (OA). Offset analgesia is defined as a disproportionately large reduction in pain perception after a small decrease in the noxious stimulation [15]. It has been described to play a crucial role in the pathological processing of pain symptoms, as it has been shown to be reduced in various chronic pain conditions (see for review [52]). Offset analgesia is based on a robust inhibitory filtering mechanism of nociceptive information using temporal pain contrast enhancement (TPCE) [4,49,62]. However, the underlying mechanisms of OA - or the TPCE phenomenon on which it is based - are not fully understood, but some research is showing that OA seems to be dominated by a complex interplay of interdependent systems at different locations along the neuroaxis, including peripheral [3,29], spinal [49] and supra-spinal mechanisms [10,28,61].

Typically, OA is tested with a tonic heat stimulus traditionally lasting for 30 to 40 seconds. The pain response of OA is often contrasted with a constant heat trial without intensity modification to control for pain adaptation, i.e., a reduction in the pain response due to a relatively long-lasting stimulus [52]. In contrast to adaptation, habituation to pain describes a reduction in pain after repeated stimulations including an interstimulus interval (ISI) [9]. In OA, the temporal contrast enhancement phenomenon has been shown with repetitive electrical stimuli [37], however a frequency of 100Hz was used which is perceived as a tonic stimulation. It remains unclear whether a continuous tonic stimulus is necessary to achieve temporal contrast enhancement of pain or the effect can also be produced via a series of clearly separated stimuli as it has been shown in other sensory modalities, e.g. in taste [47], auditory [30] and visual systems [13].

The aim of this study was to investigate whether TPCE can be observed by repetitive stimulation. It is assumed that during a repetitive TPCE stimulation, a disproportionate reduction in pain and electrodermal activity (EDA) occurs after a small reduction in heat stimuli when stimuli are clearly separated over time. In this context, it is also assumed that a shorter time interval between stimuli will show a more pronounced TPCE effect, although this effect will also be observed with longer intervals.

## 2. Methods

### Study design and procedure

This cohort study was conducted in a randomized within-subject repeated-measurement design. Following verification of eligibility criteria, oral and written information on study procedures, and informed consent, a sample of healthy pain-free participants were exposed to a heat-based repetitive TPCE paradigm using three different interstimulus intervals (ISI) in a randomized and counterbalanced sequence. Participants were seated on a chair at a table in front of a white wall during the entire procedure. All interventions and measurements were performed in an air-conditioned (temperature controlled) and quiet room. The principles of the Declaration of Helsinki were respected at all times of this study [58] and all participants could discontinue the study at any time without stating reasons. The methodology of this study was preregistered in the Open Science Framework Database (https://osf.io/4ghq3) and approved by the Ethics Committee of the University of Luebeck (21-177, 18^th^ May, 2021).

### Participants

Healthy, pain-free participants between the ages of 18 and 45 years were eligible to participate if they subjectively stated that they were healthy and pain-free on the day of examination. Participants were excluded if they had a history of chronic pain (>3 months) within the last two years, had experienced any kind of pain event (lasting longer than 30 min) within the last week and were diagnosed with neurological, cardiovascular, psychiatric or systemic diseases. Furthermore, participants were told to refrain from caffeine and cigarettes for 4 hours and from alcohol, exercise and painkillers for 24 hours before the assessment. All of them had no previous knowledge about the study aim.

### Thermal stimulation

All thermal stimuli were delivered using a Pathway CHEPS (Contact Heat -Evoked Potential Stimulator) via a 27 mm diameter contact surface (Medoc, Ramat Yishai, Israel). The CHEPS was attached to the non-dominant volar forearm (5 cm distal to the elbow). A sphygmomanometer was applied and inflated to 25 mmHg in order to standardize the contact between the skin and the thermal probe without causing pain, paresthesia or any sensory disturbances [53]. The volunteers were asked to continuously rate their pain intensity during every heat stimulus with the dominant hand on a computerized visual analogue scale (CoVAS; Medoc, Ramat Yishai, Israel) anchoring at “no pain” (=0) and “worst pain imaginable” (=100). Participants were previously familiarized with the use of the CoVAS. They were also instructed to rate pain consistently while receiving five training stimuli (familiarization: 45°C - 49°C, each 5 seconds).

For the repetitive TPCE, a protocol including two separate series of heat trials was applied: i. repetitive offset series (OS), and ii. repetitive constant series (CS). Repetitive OS included twelve consecutive heat trials, each lasting 5 seconds (s) with an ISI of 5s. The first three trials in that series were performed with an intensity of 47°C, followed by three trials with an intensity of 49°C and the final six trials with an intensity of 47°C. Repetitive CS included twelve identical heat stimuli of 47°C. The repetitive TPCE paradigm described above was administered with additional ISI of 15s and 25s. All heat stimuli were applied at identical temperature rates (rise: 70°C/s; fall: 40 °C/s). After each series, the thermal probe was moved to a new skin site to avoid interactions. A pause time of 2 minutes was applied between each series (OS, CS) and 10 minutes between each TPCE paradigm (ISI: 5s, 15s and 25s). The order of paradigms as well as the order of series within the paradigms was randomized and counterbalanced. An overview is provided in **Fig. 1**. All participants were instructed to detect small differences in pain and rate them on the CoVAS as accurately as possible.

**Fig. 1.**
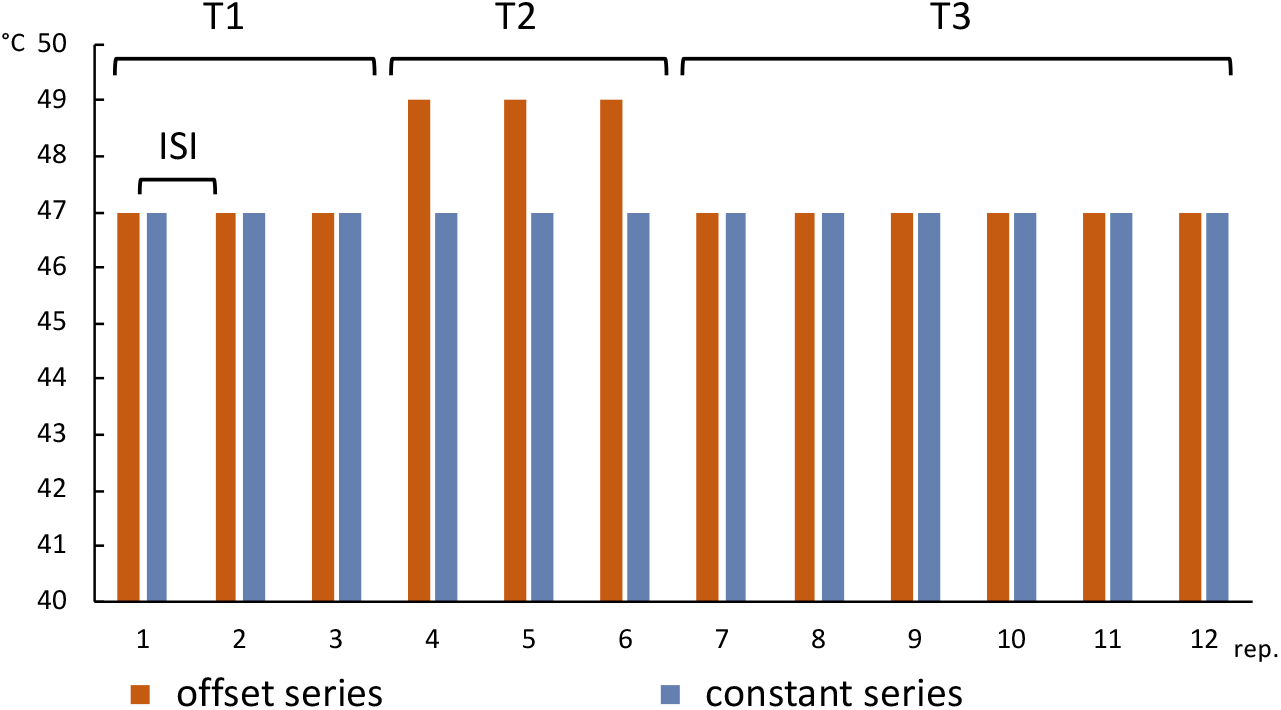
A schematic overview of the repetitive temporal pain contrast enhancement (TPCE). Two different heat stimulus series were applied: offset series (OS) and constant series (CS). Each of these series were applied with an interstimulus interval (ISI) of 5, 15 and 25 seconds in random order. The initial three heat stimuli repetitions (rep.) corresponded to the first interval (T1), the 4^th^ to 6^th^ heat stimuli formed the second interval (T2) and the remaining the third interval (T3). Each stimulus lasted 5 seconds. Temperatures of 47°C (T1 and T3 for OS and CS, as well as T2 for CS) and 49°C (T2 for OS) were contrasted.

### Electro dermal activity

During each repetitive paradigm, the respiBAN Professional (Plux, Lisbon, Portugal; a chest-worn device that registers various physiological modalities at a sampling rate of 1000 Hz) was used to collect physiological data. Electrodermal activity (EDA) signal was measured using two Ag/AgCl hydrogel electrodes (Covidien / Kendall, Dublin, Ireland) at the medial phalanx of index and middle finger of the non-dominant arm [14,40]. Electrodermal activity is dependent on sweat secretion, which is closely related to the activity in the autonomic nervous system [40]. Participants were asked to wash their hands with ordinary soap immediately before attaching the electrodes. During data collection, participants were asked not to speak and to breathe comfortably. EDA recordings was synchronized with thermal stimuli application using “opensignals” software (PLUX Wireless Biosignals, S.A., Portugal).

### Characteristics and Questionnaires

The height, weight, limb dominance and the general fear of pain measured on a 0 (=none at all) to 100 (=highest possible fear) Numerical Rating Scale (NRS) were collected before starting the heat assessment. Furthermore, the following questionnaires were administered: the Pain catastrophizing scale (PCS) includes questions about catastrophizing and ruminations to pain [22], the Patient Health Questionnaire (PHQ-9) contains questions on depression [44], the Pain and Vigilance Awareness Questionnaire (PVAQ) addresses awareness, consciousness, vigilance and pain monitoring [19], the Pain Sensitivity Questionnaire (PSQ) contains self-reported pain sensitivity based on imagined painful daily life situations [46], the Working Memory Questionnaire (WMQ) addresses three dimensions of working memory: short-term storage, attention and executive control [56].

### Data extraction and analysis

Based on the effect size for the OA response of *d* = 0.76 [15], an estimated group size of n=25 participants was required for this study (α=5%, β=95%). This was increased to n=33, in order to avoid a loss of statistical power if the previously determined effect was underestimated [1], especially given the fact that repetitive OA paradigm was not assessed previously.

Pre-processing of the CoVAS and EDA data was performed using Python Libraries. Pain rating for each millisecond was averaged to obtain one pain rating per second. All other statistical analyses were performed with the IBM Statistical Package for Social Science (SPSS Version 25, Armonk, NY). Pain ratings (CoVAS, 0-100) as well as EDA values (microsiemens, μS) were expressed as the Area Under the Curve (AUC). The time interval was chosen from the start of the corresponding heat stimulus (temperature increment from the baseline temperature) to the start of the next heat stimulus (twelve AUC values in total). This allows for a comparability between pain and EDA data, as especially the latter show a response latency. Present repetitive temporal pain contrast enhancement (ΔrTPCE) was defined as the difference between CS and OS of AUC (pain and EDA) at the 7^th^ heat stimulus (after stimulus reduction). According to the pre-registered analysis plan, this difference was analyzed using dependent two-tailed student’s t-tests (including Cohen’s d effect sizes) separately for the three different ISIs (pain: CS vs. OS, EDA: OS vs. CS). The mean AUC from the 8t^h^ to the 12^th^ stimulus was also analyzed. Also, an existing habituation effect in pain, both within the T1 interval (1^st^ vs. 3^rd^ heat stimulus), and the T2 interval (4^th^ vs. 6^th^) was examined by comparing corresponding mean values using two-tailed student’s t-tests. A repeated measure analysis of variance (rANOVA) of the ΔrTPCE response was performed including all three ISIs (5s, 15s and 25s) as within-subject factor “ISI”. Effect sizes were expressed as partial η^2^_p_. If statistically significant differences were detected, Bonferroni corrected post-hoc t-tests were performed. Pearson’s correlation (r_p_) analysis was used to evaluate the possible correlations among parametric data, whereas Spearman’s correlation analysis (r_s_) was used for not normally distributed data. Data were presented as mean and standard deviation (SD) in text and tables and as mean and standard error of the mean (SEM) in figures. The level of significance was set at p = 0.05.

## 3. Results

### 3.1. Characteristics of participants

33 healthy pain-free participants (n = 28, 85% women, n = 31, 94% right-handed) were investigated. The mean age of the subjects was 25.2 (SD 4.0) years. Further characteristics are shown in **Table 1**. No data were missing. No subject terminated the study before completion. No side effects of noxious stimulations were observed.

**Table 1.** Descriptive statistics.

| Variable | Mean (SD) |
| --- | --- |
| Height (cm) | 168.1 (28.2) |
| Weight (kg) | 68.3 (12.7) |
| Fear of pain (VAS) | 34.1 (17.3.) |
| Variable | Median (IQR) |
| PVAQ | 32.0 (28.5 - 38.0) |
| PHQ-9 | 4.0 (3.0 - 6.5) |
| PCS | 10.0 (7.5 - 13.0) |
| PSQ | 51.0 (36.0 - 65.5) |
| WMQ total | 22.0 (14.5 - 28.0) |
| storage domain | 7.0 (4.0 - 10.0) |

*IQR - interquartile range, PCS - Pain Catastrophizing Scale, PSQ - Pain Sensitivity Questionnaire, PVAQ - Pain Vigilance and Awareness Questionnaire, SD - standard deviation, VAS - Visual Analogue Scale, WMQ - Working Memory Questionnaire.*

### 3.2. Pain ratings

An illustration of the mean pain ratings per trial is shown in **Figure 2**. For all ISIs, no significant difference was found between OS and CS at the 1^st^ heat stimulus (p > 0.05) indicating that the baseline sensitivity was comparable between different series. In contrast to ISI of 25s (t_(32)_ = -1.852, p = 0.073, d = 0.32), a significant difference between OS and CS at the 7^th^ heat stimulus was found in the ISI of 5s (t_(32)_ = -5.11, **p < 0.001**, d = 0.89) and 15s (t_(32)_ = - 3.01, **p = 0.005**, d = 0.52). These results indicate that ΔrTPCE at ISI of 25s is abolished, however application of the shorter ISI preserves the ΔrTPCE effect. **Figure 3** provides an overview of the pain ratings for the OS and CS of the 7^th^ heat trial, as well as the corresponding values for the AUC for ΔrTPCE.

**Figure 2.**
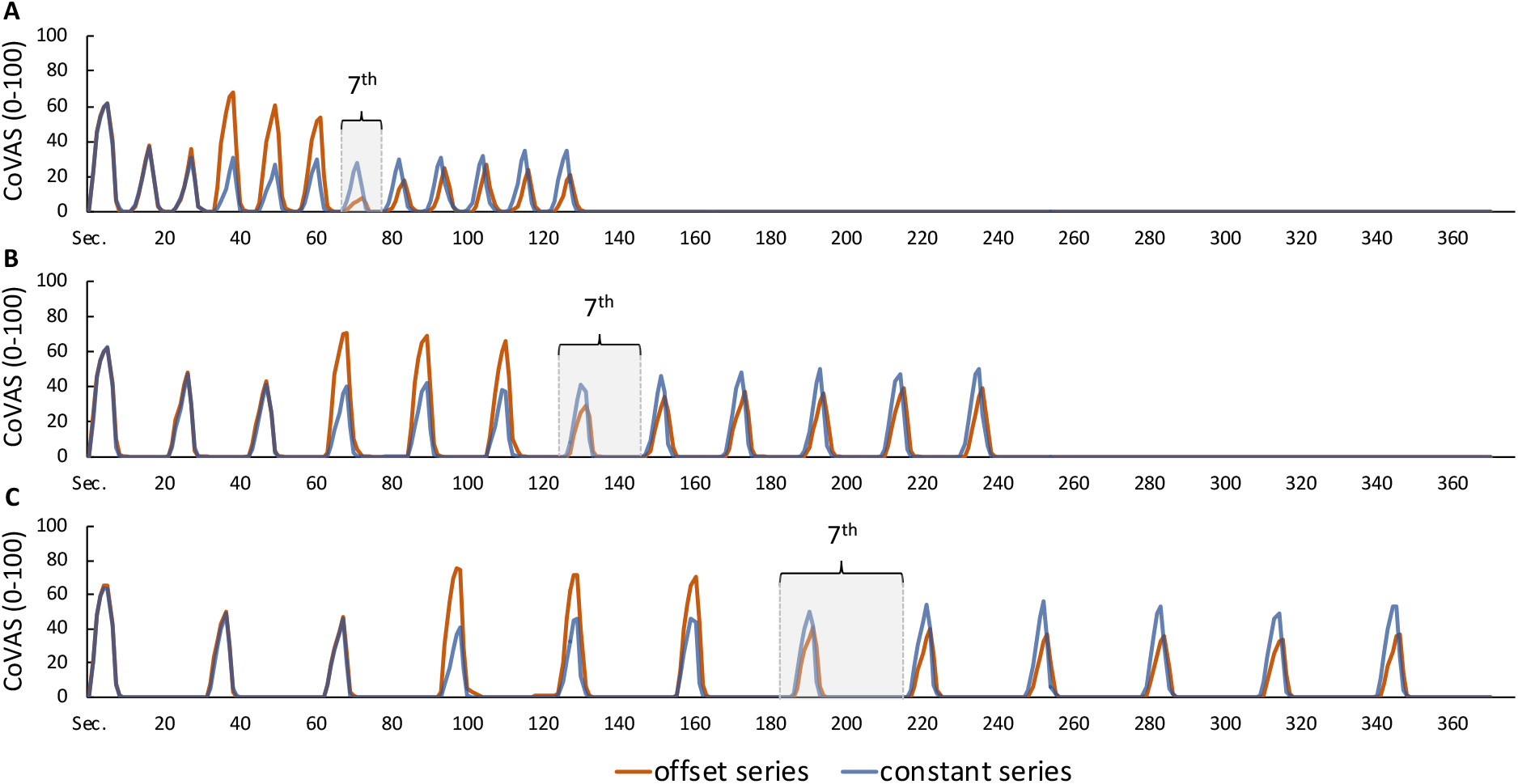
Pain ratings for all heat trials. Both the constant series (12 identical heat stimuli at a temperature of 47°C, 5 seconds each) and the offset series (3 heat stimuli at a temperature of 47°C, followed by 3 heat stimuli at 49°C and again 6 stimuli at 47°C) are presented using ratings recorded by the computerized Visual Analog Scale (CoVAS). Interstimulus intervals (ISI) of 5 (A), 15 (B) and 25 seconds (C) are compared. The pain response to the 7^th^ heat trial is indicated.

**Figure 3.**
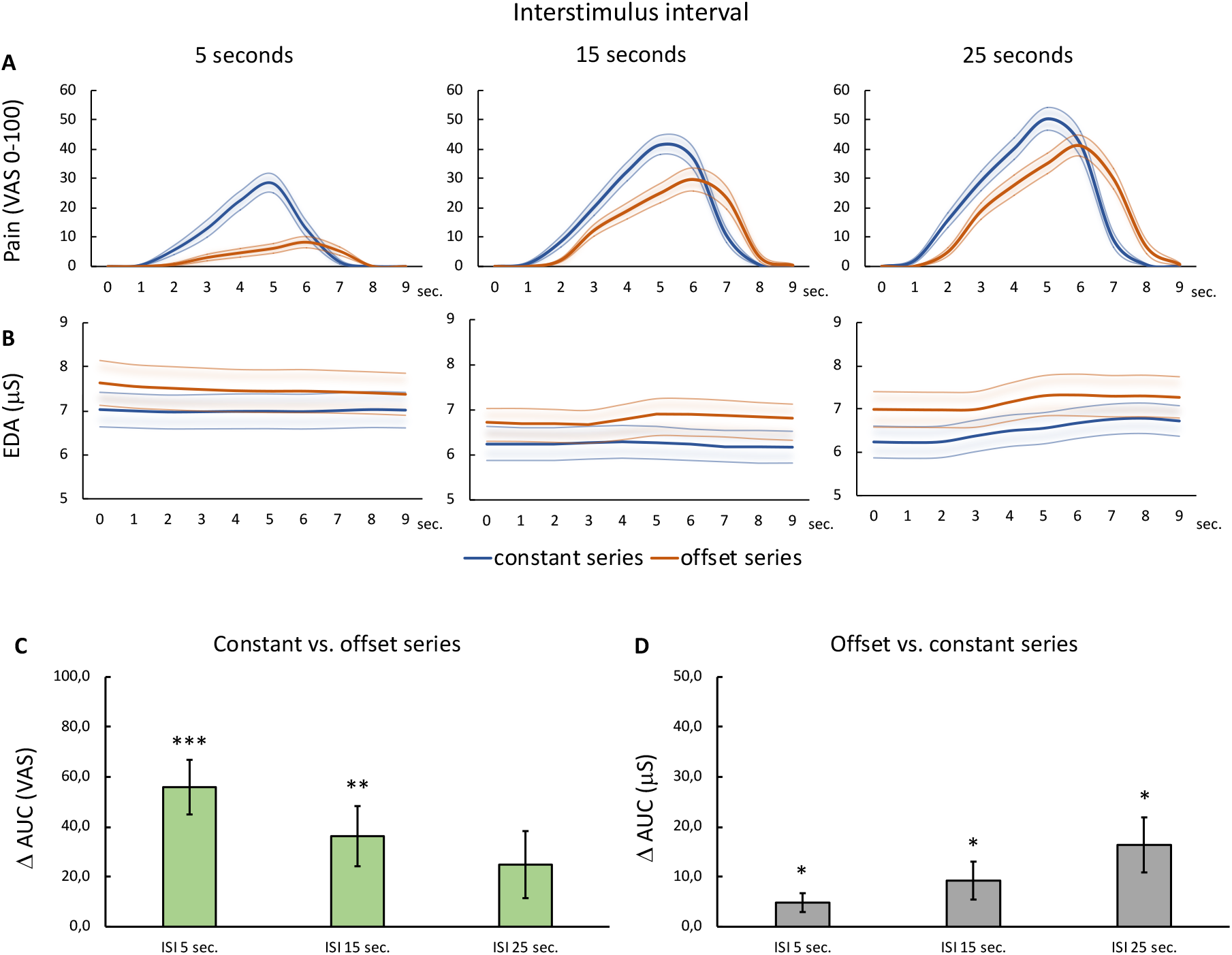
Pain ratings and electrodermal activity during the 7^th^ heat trial. In the upper and middle graphs, both the offset and constant series are shown using Computerized Visual Analog Scale (CoVAS) (A) and electrodermal activity (EDA) expressed in microsiemens (µS) (B) per interstimulus interval (ISI) 5, 15, and 25 seconds. At the bottom, the differences between the offset and the constant series for the area under the curve (AUC) for CoVAS (C) and µS (D) for each ISI are shown. All data are presented in mean values 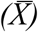 and standard error of the mean (SEM). Significant difference between OS and CS within each ISI: * p < 0.05, ** p < 0.01, *** p < 0.001. No differences were found between the ISIs.

Contrasting the average of the remaining heat trials in the T3 interval (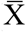 8^th^ to 12^th^) between CS and OS revealed significant differences for all ISIs (ISI of 5s: t_(32)_ = -4.13, **p < 0.001**, d = 0.72; ISI of 15s: t_(32)_ = - 3.59, **p = 0.001**, d = 0.63; ISI 25s: t_(32)_ = - 5.30, **p < 0.001**, d = 0.92). Interestingly, when the magnitude of the ΔrTPCE was compared regarding the different ISIs, no significant differences were observed. Namely, neither ΔrTPCE at the 7^th^ trial (F_(2,64)_ = 2.31, p = 0.11, 2p = 0.07) nor based on the average of the remaining trials of T3 (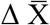 8^th^ to 12^th^) was found to be significantly different across ISIs (F_(2,64)_ = 2.40, p = 0.10, η_p_^2^ = 0.07). Additionally, 36.4% (n = 12) of participants reported having full analgesia (no pain at all) at the 7^th^ heat trial in the ISI of 5s sequence, only 3% (n = 1) at ISI 15s and none at ISI 25s. Pain was reported during all other 7^th^ heat trials in all ISI sequences. The values for the AUC of the 7^th^ trial as well as the average 8^th^ to 12^th^ trial can be found in Table 2.

**Table 2.** Pain ratings and electrodermal activity presented as the Area Under the Curve (mean, SD)

|  |  |  |  | Interstimulus interval |  |  |
| --- | --- | --- | --- | --- | --- | --- |
|  |  | Trial | Repetition | 5s | 15s | 25s |
| AUC<br>(VAS) | Pain | OS | 7<sup>th</sup> | 28.3 (39.6) | 115.2 (87.6) | 165.2 (93.6) |
|  |  | CS | 7<sup>th</sup> | 84.2 (64.7) | 151.4 (73.4) | 190.1 (92.5) |
| | | ΔrTPCE <sup>1</sup> | $\Delta$ 7<sup>th</sup> | 56.0 (62.9) | 36.3 (69.2) | 24.9 (77.1) |
| AUC<br>(μS) | EDA | OS | 7<sup>th</sup> | 81.6 (30.4) | 132.1 (50.4) | 203.7 (91.2) |
|  |  | CS | 7<sup>th</sup> | 76.8 (24.9) | 122.8 (46.1) | 187.3 (78.9) |
| | | ΔrTPCE <sup>2</sup> | $\Delta$ 7<sup>th</sup> | 4.8 (10.9) | 9.2 (21.8) | 16.4 (31.7) |
| AUC<br>(VAS) | Pain | OS | $\bar{X}$ 8<sup>th</sup> to 12<sup>th</sup> | 67.5 (53.1) | 139.8 (93.3) | 140.7 (87.2) |
| | | CS | $\bar{X}$ 8<sup>th</sup> to 12<sup>th</sup> | 102.9 (65.7) | 173.0 (77.4) | 201.4 (92.7) |
| | | ΔrTPCE <sup>1</sup> | $\Delta$ $\bar{X}$ 8<sup>th</sup> to 12<sup>th</sup> | 35.4 (49.2) | 33.2 (53.1) | 60.7 (66.1) |
| AUC<br>(μS) | EDA | OS | $\bar{X}$ 8<sup>th</sup> to 12<sup>th</sup> | 74.3 (24.9) | 123.6 (49.7) | 182.0 (82.3) |
| | | CS | $\Delta$ $\bar{X}$ 8<sup>th</sup> to 12<sup>th</sup> | 69.8 (23.4) | 117.9 (50.3) | 175.9 (77.9) |
| | | ΔrTPCE <sup>2</sup> | $\bar{X}$ 8<sup>th</sup> to 12<sup>th</sup> | -4.5 (12.2) | -5.7 (20.6) | -6.2 (29.7) |

*AUC - area under the curve, EDA - electrodermal activity, CS - constant series, OS - offset series, s - seconds, VAS - visual analogue scale (0-100), μS - microsiemens, 7<sup>th</sup> - seventh trial within the OS, CS or their difference ( $\Delta$ ), 8th to 12th presents eighth to 12th trial within the OS, CS or ( $\Delta$ ), ΔrTPCE - repetitive temporal pain contrast enhancement expressed as <sup>1</sup> the difference between CS and OS or as <sup>2</sup> the difference between OS and CS. All data are presented in mean values ( $\bar{X}$ ) and standard deviation (SD).*

Secondary exploratory analyses showed that within T1, a significant habituation occurred in all series and ISIs (T1: 1^st^ vs. 3^rd^, **p < 0.001**) and within T2 for all of the ISIs considering OS (T2: 4^th^ vs. 6^th^, **p < 0.002**), as shown by the lower pain ratings in the third stimulus compared to the first stimulus in the T1 and T2 intervals, respectively. Significant correlations between OS and CS were shown for all ISIs regarding the pain response (ISI 5s: **p = 0.035**, r_p_ = 0.368; ISI 15s: **p < 0.001**, r_p_ = 0.602; ISI 25s: **p < 0.001**, r_p_ = 0.650). Furthermore, a significant correlation was shown between the rTPCE (25s) and the PVAQ (r_s_ = -0.38, **p = 0.029**). Interestingly, the ΔrTPCE did not correlate with the total WMQ (r_s_ < 0.3, p < 0.05), but when exclusively the domain “storage” was considered, significant correlations were shown for rTPCE of 25s (r_s_ = 0.42, **p = 0.016**), but not the remaining ISI (r_s_ < 0.3, p < 0.05). No correlation was found between ΔrTPCE and other included questionnaires or characteristics (r_p,s_ < 0.3, p > 0.05).

### 3.2. Electrodermal activity

Similarly, the 1^st^ heat stimuli between OS and CS at all ISIs did not differ significantly for the EDA signal (p > 0.05). At the 7^th^ heat stimulus significant differences between CS and OS could be shown for all ISIs (5s: t_(32)_ = 2.55, **p =0.016**, d = 0.44; 15s: t_(32)_ = 2.43, **p = 0.021**, d = 0.42, 25s: t_(32)_ = 2.97, **p = 0.006**, d = 0.52). This indicates an existence of rTPCE effect at all ISIs regarding EDA (**Table 2, Figure 3**). If the average of the remaining heat trials within T3 is also compared between CS and OS (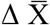 8^th^ to 12^th^), significant differences were shown exclusively for 5 seconds (t_(32)_ = 2.12, **p = 0.042**, d = 0.72), but not for the remaining ISIs (p < 0.05). As for the pain ratings, no significant differences between the three ISIs regarding the ΔrTPCE could be shown for the EDA (F_(2,64)_ = 2.19, p = 0.12, 2p = 0.06). Both the pain ratings and the corresponding EDA can be found for each heat trial (each series and ISIs) in **Figure 4**.

**Figure 4.**
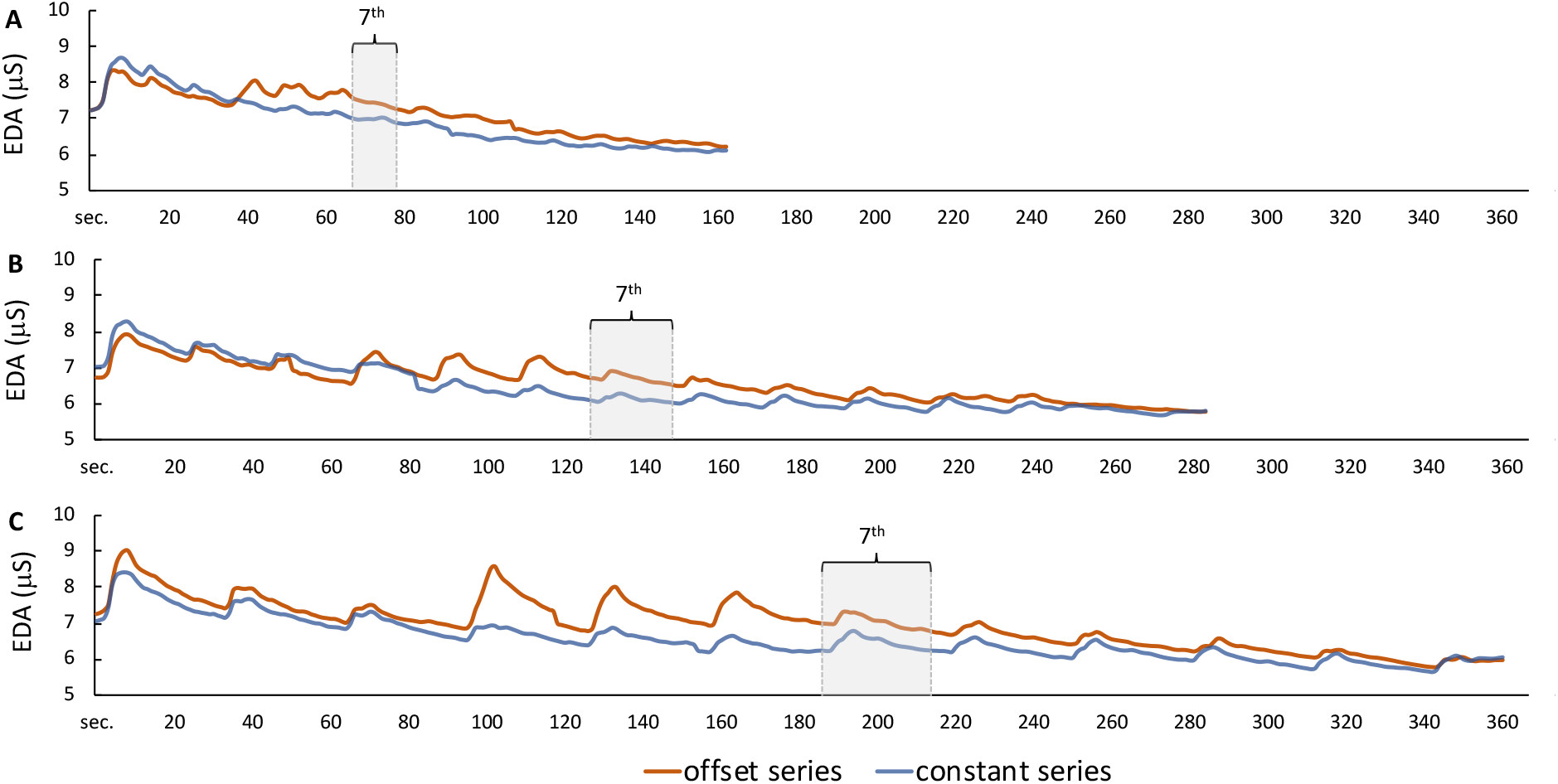
Electrodermal activity for all heat trials. Both the constant series (12 identical heat stimuli at a temperature of 47°C, 5 seconds each) and the offset series (3 heat stimuli at 47°C, followed by 3 heat stimuli at 49°C and again 6 stimuli at 47°C) are shown in electrodermal activity (EDA) expressed in microsiemens (µS). Interstimulus intervals of 5 (A), 15 (B), and 25 seconds (C) are specified. The EDA to the 7^th^ heat trial is indicated.

## 4. Discussion

Temporal contrast enhancement of pain can be induced by stimuli that are clearly separated in time with up to 25s separation. Larger periods trend to reduce TPCE, whereas - contrary to our hypothesis - no statistically significant difference between intervals were observed. Interestingly, EDA data indicate that this effect can be replicated physiologically, showing an increased EDA response - as a marker of activity in the autonomic nervous system - during temporal contrast analgesia. The EDA marker of the behavioral effect followed a reversed direction: The longer the ISI the more pronounced the difference between constant and offset series.

### Continuous temporal pain contrast enhancement

For OA, with its continuous temporal contrast enhancement of pain, again, both central and peripheral underlying mechanisms are discussed. Functional imaging studies including functional magnetic resonance imaging (fMRI) indicated the involvement of supra-spinal and spinal areas during OA [10,28,49,61,63]. Among other findings, a greater BOLD signal in the brainstem pain modulation circuit, including the periaqueductal gray (PAG) and rostral ventromedial medulla (RVM), was observed [10,61]. It is therefore particularly interesting and somewhat surprising that centrally acting drugs did not affect the OA response [21,33,34]. On the other hand, there is some evidence that specific nerve fibers are involved in the traditional continuous OA response, in which glabrous and non-glabrous skin sites were contrasted [3,29]. Likewise, an existing TPCE effect, even with repeated stimulation, may indicate central as well as peripheral mediating mechanisms.

### Pain habituation and repetitive temporal pain contrast enhancement

In general, repetitive application of noxious stimuli with an identical intensity can modulate the perceived pain intensity, which can lead to an increase (sensitization, summation) or a decrease (habituation) in reported pain [18]. There are multiple central and peripheral processes involved. One of them is temporal summation of second pain (summation via a delayed sensation of pain [41]), which involves repetitive stimulation of peripheral unmyelinated C-fibers [42]. During noxious heat, this pain summation is produced with ISIs of up to 3 seconds [32,45]. However, longer ISIs (as shown here), may lead to a reversed pain response: habituation to pain [7]. Proposed mechanisms of habituation are mainly based on central nervous system explanatory models (see [9] for a review), such as cognitive models and non-associative learning [39], and also on descending pain inhibitory activity [24]. Interestingly, habituation of pain also occurs in the long term, i.e., as a contrast of pain from day to day [5], and habituation is not limited to the context of pain but can also be observed as a behavioral response across species, stimuli and responses [43].

Peripheral mechanisms in the processing of pain habituation are also reasonable, although rather secondary. It was shown, that both A and C fiber nociceptors show fatigue responses - increased with higher stimulation - when ISIs are below 30s [48]. Thus, this afferent fatigue response may contribute to the habituation effect and one can speculate that higher stimulation (here 49°C) even potentiates this fatigue response in the 7^th^ trial (after the temperature reduction to 47°C). Furthermore, a correlation was found in the pain responses to the 7th heat trial between offset and constant series. A re-analysis of our previous OA studies [50,51,53] confirms this effect also for continuous TPCE: there is a strong correlation between the pain reduction in the offset trial and the adaptation to pain in the constant heat trial, without temperature change. Although the underlying mechanisms of adaptation to a sustained pain stimulus are not yet fully understood, again predominantly peripheral processes are hypothesized [16,57] as shown, for example, in certain A-delta nociceptors by electrophysiological studies [20,23,54,55]. However, both habituation and adaptation are strongly dependent on individual differences in pain perception with a strong tendency to inhibition. An association of habituation and the repetitive TPCE or adaptation and the continuous TPCE effect (OA) can nevertheless be hypothesized, at least in part, points to a common underlying mechanism. It should be clarified whether i) habituation and adaptation of pain are associated (our previous data suggest this), whether ii) OA and repetitive TPCE are associated, and whether iii) repetitive TPCE and OA show the same properties, such as impairment in chronic pain and unaffected by centrally acting drugs, like in OA [17].

Interesting observations made in the field of habituation point to the fact that this phenomenon can be easily disrupted. Indeed, successful habituation is impaired when repeated exposure to the same stimulus occurs through the introduction of a different or higher stimulus. This characteristic is also known as dishabituation [43]. Thus, with not just one but several higher stimuli within a habituation series (in our protocol three stimuli), the magnitude of the dishabituation produced can be expected to decrease and even become potentiated, in contrast to the original habituation. Additional associated features could be “habituation of dishabituation” and “potentiation” [9,43]. As with the mechanisms of habituation, repetitive TCPE was expected to decrease pain/irritation with more frequent stimulation over a longer period of time (or stimulation with shorter ISI). This can be explained by a temporal contrast in which the current stimulus (consciously or unconsciously) is compared and evaluated with the previous stimulus. It is supported by the present association with short-term storage in working memory, which corresponds to the ability to retain information in short-term memory, and with the association of attention, vigilance and observation of pain (PAVQ), here at least for longer ISIs. For this, a trend but no significant difference between the presented ISIs could be shown. Reasons for this could be a high variance of the inter-individual pain responses [11]. Intentionally, no individualization was chosen in this experiment to allow a better relation between pain and the autonomic nervous system as this was shown to be more associated with stimulus intensity than with pain [31]. Further studies with an individualized stimulus are required to confirm this.

### Autonomic function and repetitive temporal pain contrast enhancement

A behavioral effect described by TPCE was successfully shown by the physiological effect measured via EDA. A significantly higher autonomic arousal was observed after higher stimulus intensity (here 49 °C) compared to the condition wherein each stimulus had the same intensity of 47 degrees. This observation may indicate that TPCE has a central component driven by top-down processes likely originated in the PAG. Again, this brain region has been shown to be involved in conventional continuous TPCE (OA) [10,61] and seems to have a central role in both pain inhibition and (at least in part) in autonomic response [26,27]. Our data, however, indicate that the relationship between the contrast effect and autonomic arousal is inverse. Paradoxically, less pain ‘means’ more arousal. It is not surprising that a greater effect size (d= 0.52) was observed for longer stimuli delay. EDA reflects local changes in skin conduction that depend on sweat gland secretion. Typically, the half-recovery period is between 2 to 10s explaining why longer ISIs led to greater physiological contrast [8].

One can speculate that enhanced EDA responses in post-stimulus offset can serve as inhibitory markers. It can be assumed that the higher temperature (49°C) in trials four to six could serve as the conditioning stimulus that inhibits subsequent pain originated from trials seventh to twelve. This might be the local inhibition resembling pain-inhibiting-pain effect [35], in which remote stimulus of a higher intensity can trigger descending pain control lasting for as long as 5 minutes [12,25]. Whether indeed TPCE resembles the pain-inhibiting-pain effect must be determined in future experiments with longer ISIs.

### Clinical implication

The investigation of repetitive TPCE may contribute to a better understanding of the descending pain modulation systems. Thus, it can be hypothesized that the underlying mechanisms of repetitive noxious stimulation leading to habituation to pain partially overlaps with the pain response of repetitive TPCE. This assumption is based on the repetitive nature, contrast to habituation, and anti-nociceptive properties of these two paradigms. However, a direct comparison between continuous and repetitive paradigms was not the subject of this work.

A large number of studies have described a possible relationship between preoperative quantitative sensory testing procedures and chronic postoperative pain (see for review [38]). In particular, it has been shown that descending pain modulation quantification procedures, such as temporal pain summation and pain inhibiting pain effect, have the most consistent predictive values for chronic postoperative pain and analgesic effect. Accordingly, in case of (dys)functionality, different and more effective therapeutic pathways should be offered in each individual clinical setting. Other testing procedures such as OA have not been studied in this regard, although different test paradigms seem to reflect different sub-aspects of pain modulation [53] and this might allow better phenotyping [59,60]. One reason for this could be that procedures such as OA are laborious procedures that require expensive equipment. This project, on the other hand, helps to provide the basis for easier and less expensive variants that could be applied within in a bedside manner using, for example, pressure algometry.

## Acknowledgments

The authors have no conflicts of interest to declare. There was no financial support for this project. The authors thank the Institute of Medical Informatics, University of Luebeck, kindly for providing the research facilities and equipment.

## References

[1] Anderson CJ. Central Limit Theorem. The Corsini Encyclopedia of Psychology. American Cancer Society, 2010. pp. 1–2. doi:10.1002/9780470479216.corpsy0160.

[2] Arendt-Nielsen L. Central Sensitization in Humans: Assessment and Pharmacology. In: Schaible H-G, editor. Pain Control. Handbook of Experimental Pharmacology. Berlin, Heidelberg: Springer, 2015. pp. 79–102. doi:10.1007/978-3-662-46450-2_5.

[3] Asplund CL, Kannangath A, Long VJE, Derbyshire SWG. Offset analgesia is reduced on the palm and increases with stimulus duration. Eur J Pain 2021;25:790–800.

[4] Beck B, Gnanasampanthan S, Iannetti GD, Haggard P. No temporal contrast enhancement of simple decreases in noxious heat. J Neurophysiol 2019;121:1778–1786.

[5] Bingel U, Schoell E, Herken W, Büchel C, May A. Habituation to painful stimulation involves the antinociceptive system. Pain 2007;131:21–30.

[6] Bingel U, Tracey I. Imaging CNS modulation of pain in humans. Physiology (Bethesda) 2008;23:371–380.

[7] Condes-Lara M, Calvo JM, Fernandez-Guardiola A. Habituation to bearable experimental pain elicited by tooth pulp electrical stimulation. Pain 1981;11:185–200.

[8] Dawson ME, Schell AM, Filion DL. The Electrodermal System. In: Berntson GG, Cacioppo JT, Tassinary LG, editors. Handbook of Psychophysiology. Cambridge Handbooks in Psychology. Cambridge: Cambridge University Press, 2016. pp. 217–243. doi:10.1017/9781107415782.010.

[9] De Paepe AL, Williams AC de C, Crombez G. Habituation to pain: a motivational-ethological perspective. Pain 2019;160:1693–1697.

[10] Derbyshire SWG, Osborn J. Offset analgesia is mediated by activation in the region of the periaqueductal grey and rostral ventromedial medulla. Neuroimage 2009;47:1002–1006.

[11] Fillingim RB. Individual differences in pain: understanding the mosaic that makes pain personal. Pain 2017;158 Suppl 1:S11–S18.

[12] France CR, Suchowiecki S. A comparison of diffuse noxious inhibitory controls in men and women. Pain 1999;81:77–84.

[13] Georgeson MA. Temporal properties of spatial contrast vision. Vision Res 1987;27:765–780.

[14] Gouverneur P, Li F, Adamczyk WM, Szikszay TM, Luedtke K, Grzegorzek M. Comparison of Feature Extraction Methods for Physiological Signals for Heat-Based Pain Recognition. Sensors (Basel) 2021;21:4838.

[15] Grill JD, Coghill RC. Transient analgesia evoked by noxious stimulus offset. J Neurophysiol 2002;87:2205–2208.

[16] Hashmi JA, Davis KD. Effects of temperature on heat pain adaptation and habituation in men and women. Pain 2010;151:737–743.

[17] Hermans L, Calders P, Van Oosterwijck J, Verschelde E, Bertel E, Meeus M. An Overview of Offset Analgesia and the Comparison with Conditioned Pain Modulation: A Systematic Literature Review. Pain Physician 2016;19:307–326.

[18] Jepma M, Jones M, Wager TD. The dynamics of pain: evidence for simultaneous site-specific habituation and site-nonspecific sensitization in thermal pain. J Pain 2014;15:734–746.

[19] Kunz M, Capito ES, Horn-Hofmann C, Baum C, Scheel J, Karmann AJ, Priebe JA, Lautenbacher S. Psychometric Properties of the German Version of the Pain Vigilance and Awareness Questionnaire (PVAQ) in Pain-Free Samples and Samples with Acute and Chronic Pain. IntJ Behav Med 2017;24:260–271.

[20] LaMotte RH, Thalhammer JG, Robinson CJ. Peripheral neural correlates of magnitude of cutaneous pain and hyperalgesia: a comparison of neural events in monkey with sensory judgments in human. J Neurophysiol 1983;50:1–26.

[21] Martucci KT, Eisenach JC, Tong C, Coghill RC. Opioid-independent mechanisms supporting offset analgesia and temporal sharpening of nociceptive information. Pain 2012;153:1232–1243.

[22] Meyer K, Sprott H, Mannion AF. Cross-cultural adaptation, reliability, and validity of the German version of the Pain Catastrophizing Scale. Journal of Psychosomatic Research 2008;64:469–478.

[23] Meyer RA, Campbell JN. Evidence for two distinct classes of unmyelinated nociceptive afferents in monkey. Brain Res 1981;224:149–152.

[24] Millan MJ. Descending control of pain. Prog Neurobiol 2002;66:355–474.

[25] Motohashi K, Umino M. Heterotopic painful stimulation decreases the late component of somatosensory evoked potentials induced by electrical tooth stimulation. Brain Res Cogn Brain Res 2001;11:39–46.

[26] Nahman-Averbuch H, Dayan L, Sprecher E, Hochberg U, Brill S, Yarnitsky D, Jacob G. Pain Modulation and Autonomic Function: The Effect of Clonidine. Pain Med 2016;17:1292–1301.

[27] Nahman-Averbuch H, Dayan L, Sprecher E, Hochberg U, Brill S, Yarnitsky D, Jacob G. Sex differences in the relationships between parasympathetic activity and pain modulation. Physiol Behav 2016;154:40–48.

[28] Nahman-Averbuch H, Martucci KT, Granovsky Y, Weissman-Fogel I, Yarnitsky D, Coghill RC. Distinct brain mechanisms support spatial vs temporal filtering of nociceptive information. Pain 2014;155:2491–2501.

[29] Naugle KM, Cruz-Almeida Y, Fillingim RB, Riley JL. Offset analgesia is reduced in older adults. Pain 2013;154:2381–2387.

[30] Needham JG. Contrast effects in judgments of auditory intensities. Journal of Experimental Psychology 1935;18:214–226.

[31] Nickel MM, May ES, Tiemann L, Postorino M, Ta Dinh S, Ploner M. Autonomic responses to tonic pain are more closely related to stimulus intensity than to pain intensity. Pain 2017;158:2129–2136.

[32] Nielsen null, Arendt-Nielsen null. The importance of stimulus configuration for temporal summation of first and second pain to repeated heat stimuli. Eur J Pain 1998;2:329– 341.

[33] Niesters M, Dahan A, Swartjes M, Noppers I, Fillingim RB, Aarts L, Sarton EY. Effect of ketamine on endogenous pain modulation in healthy volunteers. Pain 2011;152:656–663.

[34] Niesters M, Hoitsma E, Sarton E, Aarts L, Dahan A. Offset analgesia in neuropathic pain patients and effect of treatment with morphine and ketamine. Anesthesiology 2011;115:1063–1071.

[35] Nir R-R, Yarnitsky D. Conditioned pain modulation. Curr Opin Support Palliat Care 2015;9:131–137.

[36] Ossipov MH, Morimura K, Porreca F. Descending pain modulation and chronification of pain. Curr Opin Support Palliat Care 2014;8:143–151.

[37] Petersen KK, Mørch CD, Ligato D, Arendt-Nielsen L. Electrical stimulation for evoking offset analgesia: A human volunteer methodological study. Eur J Pain 2018;22:1678–1684.

[38] Petersen KK, Vaegter HB, Stubhaug A, Wolff A, Scammell BE, Arendt-Nielsen L, Larsen DB. The predictive value of quantitative sensory testing: a systematic review on chronic postoperative pain and the analgesic effect of pharmacological therapies in patients with chronic pain. Pain 2021;162:31–44.

[39] Poon C-S, Schmid S. Nonassociative Learning. In:Seel NM, editor. Encyclopedia of the Sciences of Learning. Boston, MA: Springer US, 2012. pp. 2475–2477. doi:10.1007/978-1-4419-1428-6_1849.

[40] Posada-Quintero HF, Chon KH. Innovations in Electrodermal Activity Data Collection and Signal Processing: A Systematic Review. Sensors (Basel) 2020;20:E479.

[41] Price DD. Characteristics of second pain and flexion reflexes indicative of prolonged central summation. Exp Neurol 1972;37:371–387.

[42] Price DD, Hu JW, Dubner R, Gracely RH. Peripheral suppression of first pain and central summation of second pain evoked by noxious heat pulses. Pain 1977;3:57–68.

[43] Rankin CH, Abrams T, Barry RJ, Bhatnagar S, Clayton D, Colombo J, Coppola G, Geyer MA, Glanzman DL, Marsland S, McSweeney F, Wilson DA, Wu C-F, Thompson RF. Habituation Revisited: An Updated and Revised Description of the Behavioral Characteristics of Habituation. Neurobiol Learn Mem 2009;92:135–138.

[44] Reich H, Rief W, Brähler E, Mewes R. Cross-cultural validation of the German and Turkish versions of the PHQ-9: an IRT approach. BMC Psychology 2018;6:26.

[45] Riley JL, Cruz-Almeida Y, Staud R, Fillingim RB. Effects of manipulating the interstimulus interval on heat-evoked temporal summation of second pain across the age span. Pain 2019;160:95–101.

[46] Ruscheweyh R, Marziniak M, Stumpenhorst F, Reinholz J, Knecht S. Pain sensitivity can be assessed by self-rating: Development and validation of the Pain Sensitivity Questionnaire. Pain 2009;146:65–74.

[47] Schifferstein HN, Oudejans IM. Determinants of cumulative successive contrast in saltiness intensity judgments. Percept Psychophys 1996;58:713–724.

[48] Slugg RM, Meyer RA, Campbell JN. Response of cutaneous A-and C-fiber nociceptors in the monkey to controlled-force stimuli. J Neurophysiol 2000;83:2179–2191.

[49] Sprenger C, Stenmans P, Tinnermann A, Büchel C. Evidence for a spinal involvement in temporal pain contrast enhancement. Neuroimage 2018;183:788–799.

[50] Szikszay TM, Adamczyk WM, Carvalho GF, May A, Luedtke K. Offset analgesia: somatotopic endogenous pain modulation in migraine. Pain 2019.

[51] Szikszay TM, Adamczyk WM, Hoegner A, Woermann N, Luedtke K. The effect of acute-experimental pain models on offset analgesia. Eur J Pain 2021;25:1150–1161.

[52] Szikszay TM, Adamczyk WM, Luedtke K. The Magnitude of Offset Analgesia as a Measure of Endogenous Pain Modulation in Healthy Participants and Patients With Chronic Pain: A Systematic Review and Meta-Analysis. Clin J Pain 2019;35:189–204.

[53] Szikszay TM, Lévénez JLM, von Selle J, Adamczyk WM, Luedtke K. Investigation of correlations between pain modulation paradigms. Pain Med 2021.

[54] Treede RD, Meyer RA, Campbell JN. Myelinated mechanically insensitive afferents from monkey hairy skin: heat-response properties. J Neurophysiol 1998;80:1082–1093.

[55] Treede RD, Meyer RA, Raja SN, Campbell JN. Evidence for two different heat transduction mechanisms in nociceptive primary afferents innervating monkey skin. J Physiol 1995;483 (Pt 3):747–758.

[56] Vallat-Azouvi C, Pradat-Diehl P, Azouvi P. The Working Memory Questionnaire: A scale to assess everyday life problems related to deficits of working memory in brain injured patients. Neuropsychological Rehabilitation 2012;22:634–649.

[57] Weissman-Fogel I, Dror A, Defrin R. Temporal and spatial aspects of experimental tonic pain: Understanding pain adaptation and intensification. Eur J Pain 2015;19:408–418.

[58] World Medical Association. World Medical Association Declaration of Helsinki: ethical principles for medical research involving human subjects. JAMA 2013;310:2191–2194.

[59] Yarnitsky D. Role of endogenous pain modulation in chronic pain mechanisms and treatment. Pain2015;156 Suppl 1:S24–31.

[60] Yarnitsky D, Granot M, Granovsky Y. Pain modulation profile and pain therapy: between pro-and antinociception. Pain 2014;155:663–665.

[61] Yelle MD, Oshiro Y, Kraft RA, Coghill RC. Temporal filtering of nociceptive information by dynamic activation of endogenous pain modulatory systems. J Neurosci 2009;29:10264–10271.

[62] Yelle MD, Rogers JM, Coghill RC. Offset analgesia: a temporal contrast mechanism for nociceptive information. Pain 2008;134:174–186.

[63] Zhang S, Li T, Kobinata H, Ikeda E, Ota T, Kurata J. Attenuation of offset analgesia is associated with suppression of descending pain modulatory and reward systems in patients with chronic pain. Mol Pain 2018;14:1744806918767512.

